# Association of ulcerative colitis with atopic dermatitis: identification of shared and unique mechanisms by construction and computational analysis of disease maps

**DOI:** 10.1101/2025.04.23.650149

**Authors:** Oxana Lopata, Marcio Luis Acencio, Xinhui Wang, Ahmed Abdelmonem Hemedan, Michael J. Chao, Scott A Jelinsky, Florian Tran, Philip Rosenstiel, Andrew Y.F. Li Yim, Reinhard Schneider, Venkata Satagopam, Marek Ostaszewski

## Abstract

**Background and Aims:** Ulcerative colitis (UC) and atopic dermatitis (AD) are immune-mediated inflammatory diseases with limited treatment options. They are known to be related which may explain higher risk of development of UC in patients with AD. The goal of this work is to review and analyse molecular mechanisms of UC in comparison to AD towards insights into UC complexity, potential comorbidities and novel therapies.

**Methods:** We developed graphical computational models of UC and AD molecular mechanisms (disease maps) by integrating information from over 800 manually curated articles. The maps are available online at https://imi-immuniverse.elixir-luxembourg.org. Disease-specific risk variants and gene expression profiles are visualised to identify signatures specific to UC, and shared with AD. Computational analysis shows key proteins, their interactions and pathways shared between UC and AD.

**Results:** UC and AD maps include more than 2000 molecular interactions. Systematic computational comparison shows that both disorders exhibit epithelial barrier dysfunction, immune dysregulation involving abnormal Th2, Th1 and ILC response, common inflammatory pathways and biomarkers such as IL-13, IL-4R, IFNG, and IL-18. Visualisation and analysis of omics data demonstrates UC map usability.

**Conclusions:** We developed the first computational graphic model of UC molecular mechanisms. Its content focuses on mechanisms of epithelial barrier disruption and downstream immune dysregulation through cytokines and other mediators. The comparison of UC to AD mechanisms demonstrates common signalling pathways and biomarkers, and supports potential drug repurposing and treatment options. The workflow can be reused and new findings can be dynamically integrated into the maps.

GRAPHICAL ABSTRACT

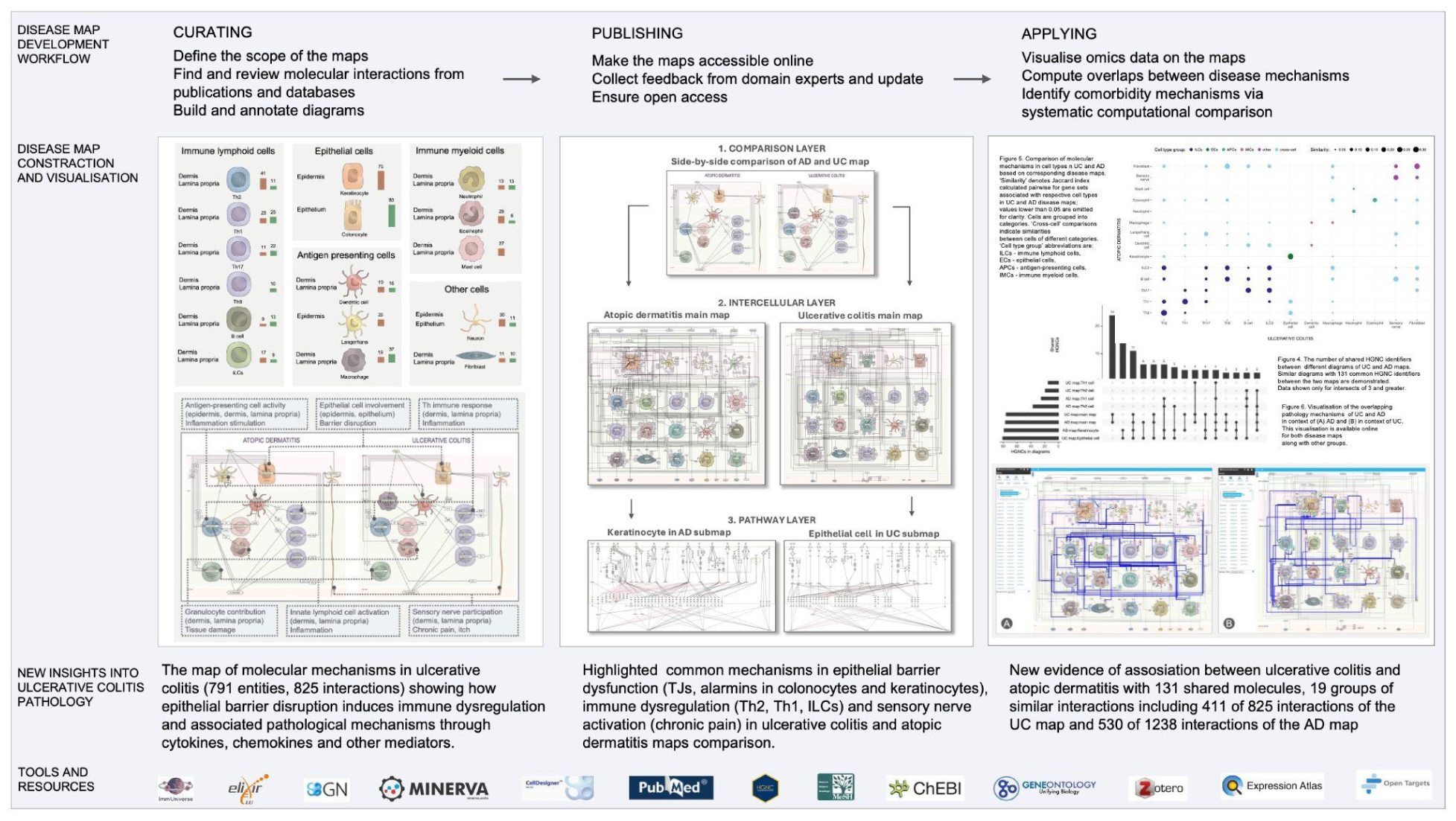

## 1. INTRODUCTION

Immune-mediated inflammatory diseases (IMIDs) are a group of disorders characterised by dysregulation of the immune system causing persistent inflammation and tissue damage. IMIDs cause physical pain and discomfort, limiting functional abilities, and reducing overall quality of life, and may lead to premature death^1,2^. A significant number of patients either do not respond to treatment or experience a recurrence of their condition after initially successful therapy, with the risk of progression of subsequent IMIDs^3^.

Various conditions can arise that manifest into different organs and tissues, such as inflammatory arthritis (IA), inflammatory bowel disease (IBD), and skin-related inflammatory disorders. Recent research suggests a potential link between chronic skin diseases and gastrointestinal disorders^4,5^. Ulcerative colitis (UC) and atopic dermatitis (AD) are both IMIDs that share common etiological factors. UC is a chronic inflammatory bowel disease that affects the colon and is characterised by recurring inflammation and ulcers of the colon’s mucosal lining, starting in the rectum and extending upwards^6^. AD is a recurrent, chronic inflammatory skin disease characterised by an impaired epidermal barrier, severe skin inflammation, cutaneous infections, and pruritus^7^. Both UC and AD involve dysfunctions in barrier integrity, UC with the intestinal lining and AD with the skin, that contribute to their pathogenesis^8,9^.

AD is associated with a number of comorbidities, including UC^4^. AD influenced patients with IBD and especially with UC^10^. High risk of development of UC in patients with AD was shown in retrospective UK and German cohort analysis^11,12^, in the Japanese^13^ and Korean^14^ populations, and meta-analysis of cross-disease relationship demonstrates that patients with AD have an increased risk of developing UC^5^. Moreover, genetic analyses support the UC-AD link, notably their causal relationships using Mendelian randomization (MR) studies in European^15^ and East Asian^16^ AD populations.

These connections suggest shared pathological molecular pathways which contribute to the development of UC and AD. However, available resources lack the capacity to study such pathways in these complex conditions. In this work we present the UC and AD disease maps, computational and diagrammatic models of molecular mechanisms implicated in both disorders, following systems biology standards, based on relevant biomedical research data. The maps are developed on systematic curation of relevant literature and constructed using established systems biomedicine methods^17,18^. We further refined the maps using omics databases to ensure their relevance. These maps serve as an interactive graphical tool that integrates disease mechanisms and reviews pathological pathways. This standardised knowledge resource is a collection of interconnected interactive diagrams following systems biology standards. Biocuration and diagram construction follow established guidelines^19^, and use community standards and workflows (https://disease-maps.io). Our maps are an open access online resource, allowing interactive exploration^1^. Finally, the maps support computational workflows, network analysis, or modelling approaches.

In this article, we outline the design of the UC and AD disease maps, their hierarchy and connectivity. We describe methods we used for map development, encoding interactions into diagrams, and annotation map elements, analyse disease phenotypes, identify common and disease-specific mechanisms and highlight shared aspects between UC and AD. We demonstrate the application of the UC map by integrating and interpreting colitis genetic risk factors from the OpenTargets platform and expression profiles from ExpressionAtlas. Finally, we programmatically compare UC and AD disease maps identifying shared molecules, cell types and molecular network modules.

## 2. RESULTS

The UC map is a state-of-the-art resource, which includes information from 325 disease-relevant publications with 791 entities, 825 interactions and 327 unique molecular entities (proteins, RNAs, genes and simple molecules), every one confirmed with a reference to a publication with experimentally validated evidence. The second large part of the project is the AD map, based on 478 publications, with 1087 entities, 1251 interactions and 391 unique molecular entities.

The UC and AD disease maps are available as an open access, interactive online resource at https://imi-immuniverse.elixir-luxembourg.org. The user guide and licensing information are available online at https://disease-maps.io/ucadmap.

Below, we describe the design of UC and AD maps, outline the contents of their diagrams and present use cases for their application.

### 2.1 Structure of UC and AD maps

The maps represent knowledge from publications about UC and AD molecular and cellular mechanisms. The contents of the maps were developed based on manual curation of scientific publications guided by focused literature search, clinical expertise and refined by computational analysis. We encoded this information on the level of intra- and intercellular interactions in a comprehensive and systematic way following an established protocol^19^.

UC and AD disease maps include three layers (Figure 1):

1. **Comparison layer** - is an overview diagram in a compact and illustrative side-by-side representation of key cell types, cytokines and signalling pathways. This overview diagram is a simplified graphical representation of disease mechanisms, meant as an entry point for detailed UC and AD maps.
2. **Intercellular layer** - represents a network of key cell types and molecular interactions between them. These diagrams contain signalling interactions and follow systems biology standards and are human- and computer-readable.
3. **Pathway layer** - describes disease and cell type specific pathways composed of metabolic, signalling and gene regulatory interactions. They follow systems biology standards and are human- and computer-readable.

**Figure 1.**
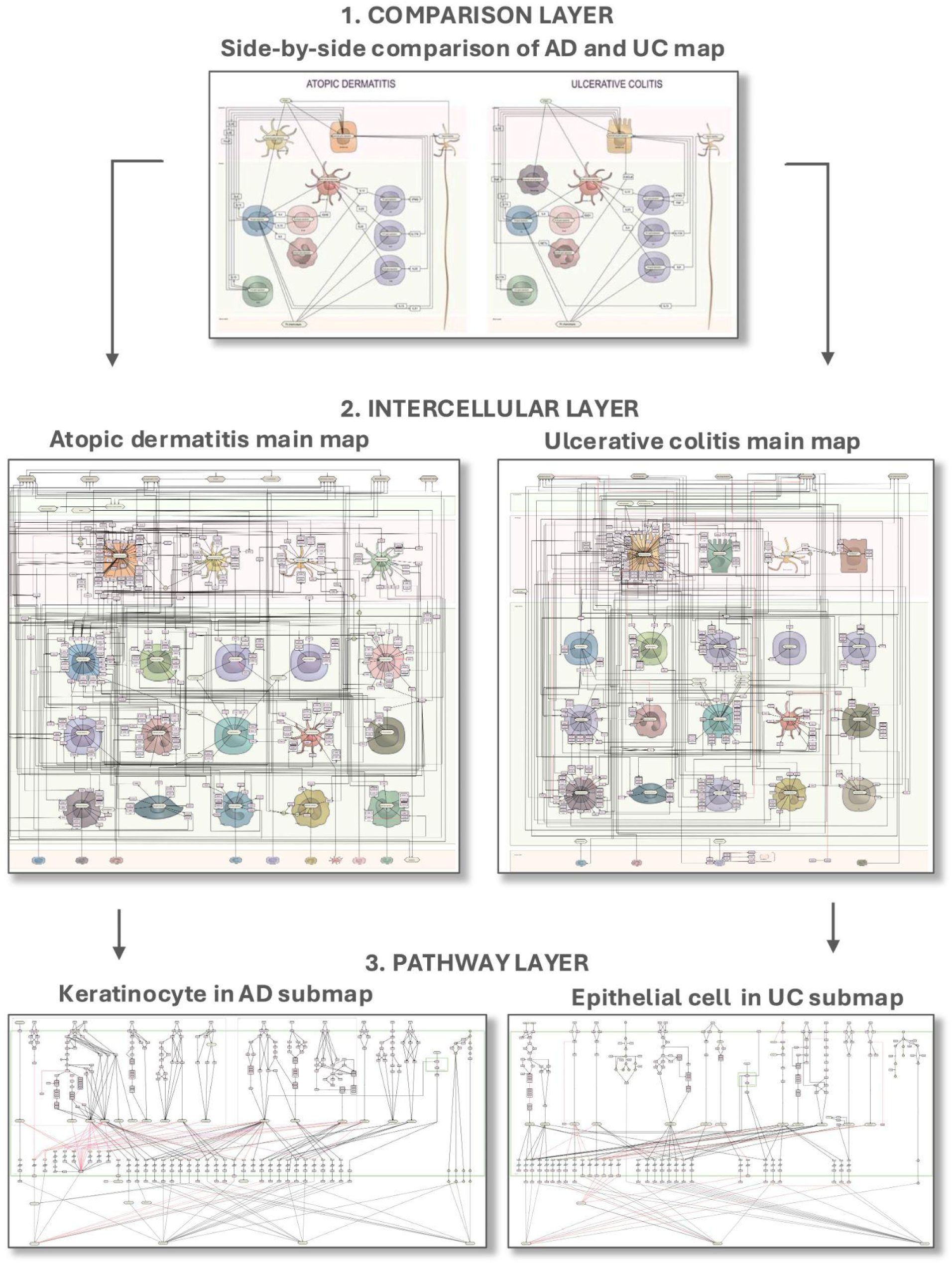
Schematic review of the UC and AD disease maps. The first layer is a compact illustrative side-by-side comparison of UC and AD mechanisms. The second layer shows individual maps of UC and AD with cells and interactions between them through cytokines, chemokines, simple molecules and the corresponding receptors. The third layer describes specific pathways in key cell types for each disease. The keratinocyte diagram for AD and the epithelial cell diagram for UC are shown as an example. The user guide is available online at https://disease-maps.io/ucadmap.

### 2.2 Comparison layer: side-by-side representation of key UC and AD mechanisms

UC and AD are distinct conditions with their own unique characteristics and manifestations. At the same time they share common underlying mechanisms.

While the UC and AD pathology maps were built independently, certain mechanism-related features appeared important for both diseases. The disease comparison layer summarises these aspects for both diseases (Figure 2).

**Figure 2.**
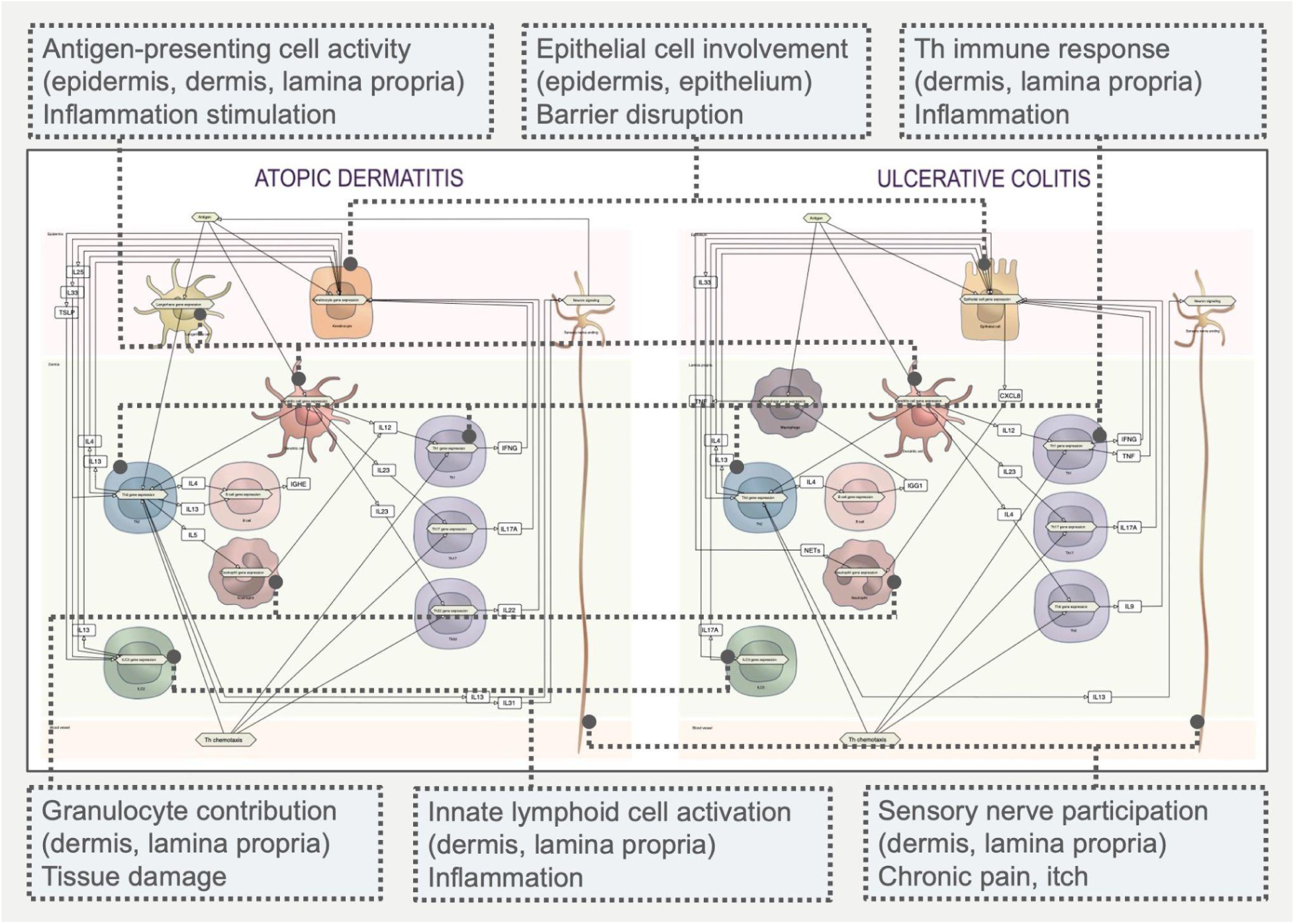
UC and AD side-by-side comparison represents common mechanisms of these two conditions. Similar cell types and their involvement in disease patterns are shown with boxes and corresponding connectors.

**Figure 3.**
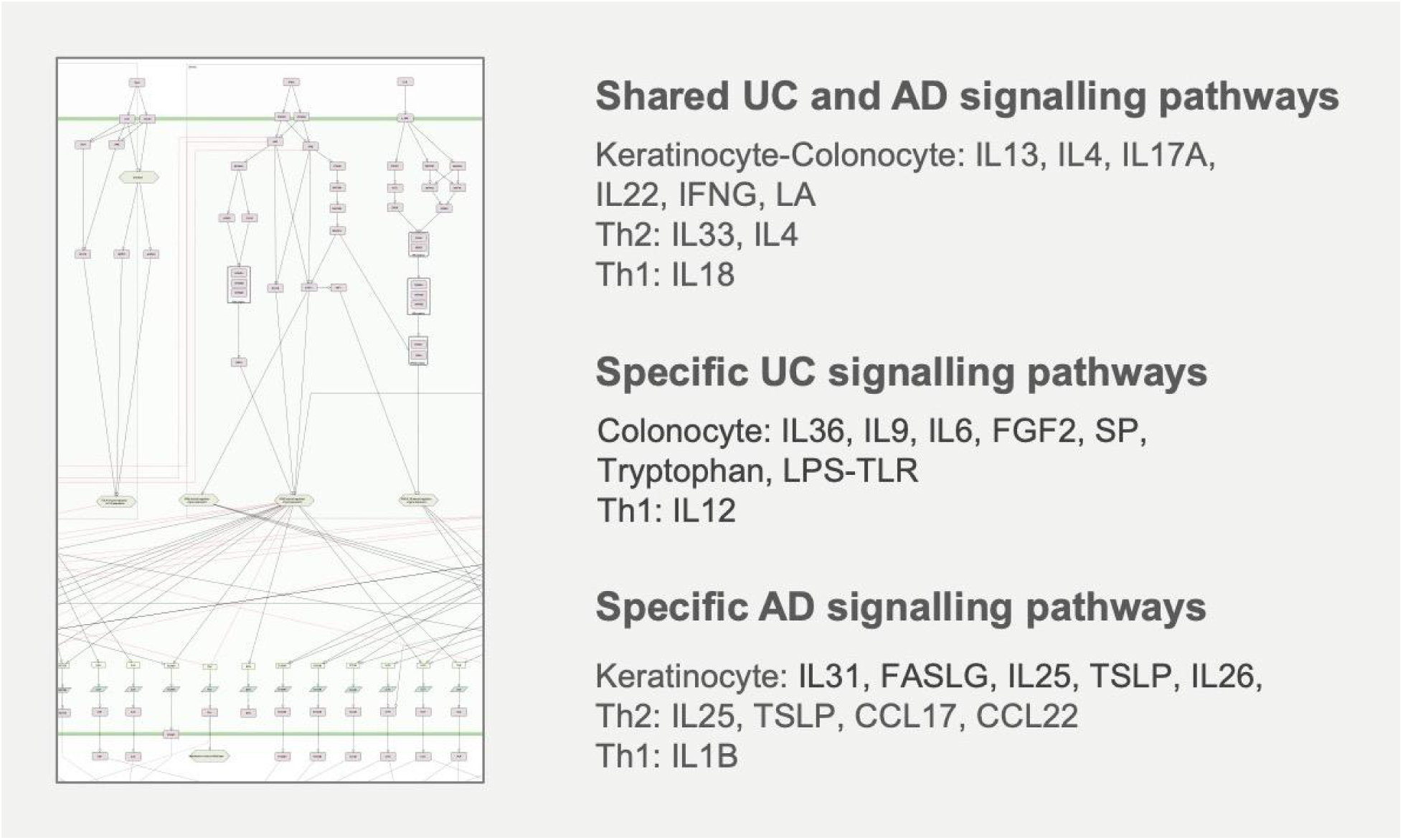
Intracellular mechanisms in the pathway layer. Common and unique signaling pathways in submaps of key cell types for UC and AD are listed.

**Figure 4.**
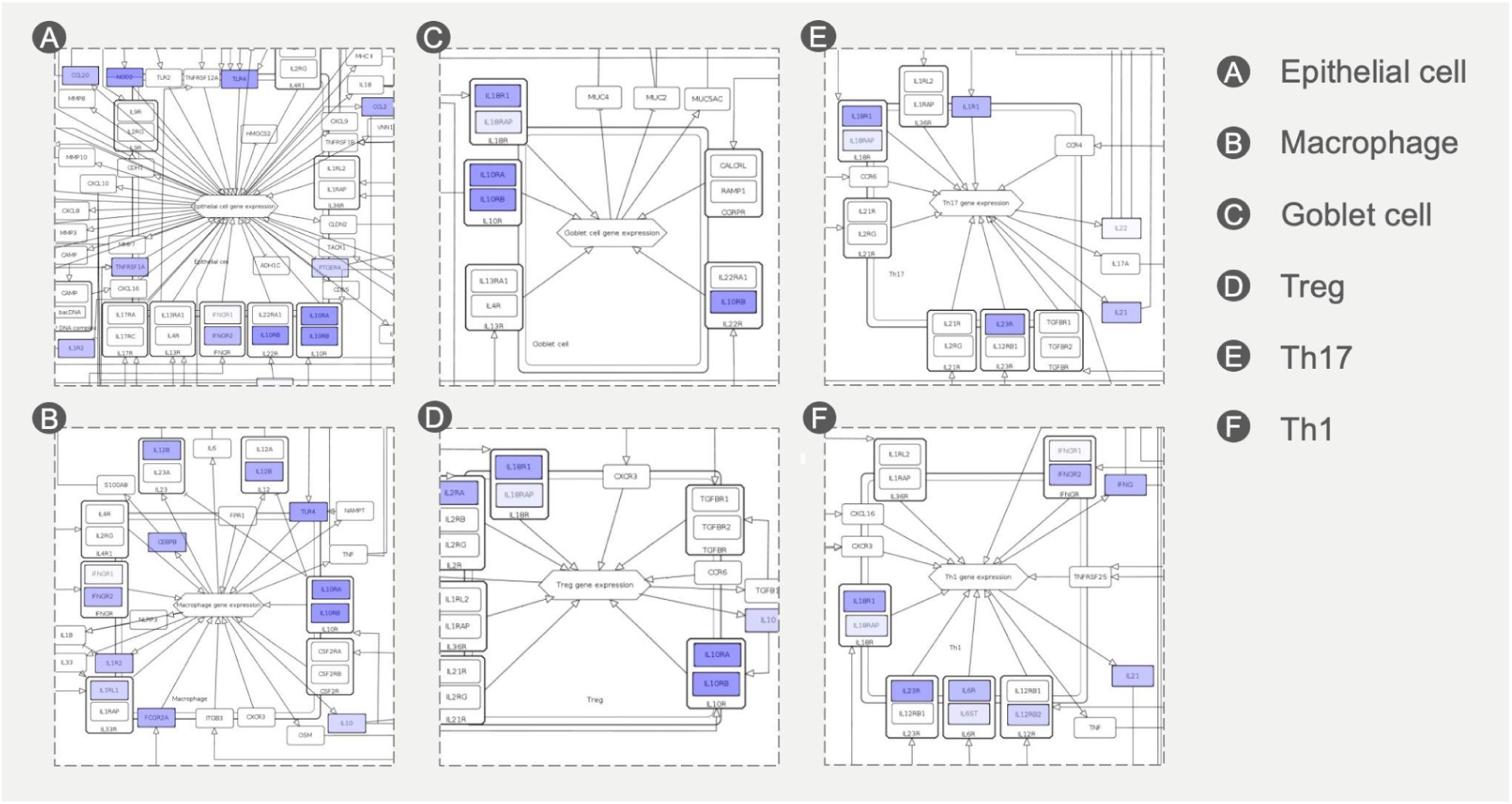
Visualization of UC-associated gene variants across different cell types. UC-linked SNPs, highlighted in blue, identified from the Open Targets Genetics, are represented in different immune and epithelial cell types in the UC disease map.

The comparison map demonstrates the following common mechanisms:

- involvement of epithelial cells - keratinocytes and colonocytes - in barrier disruption and triggering inflammation;
- antigen-presenting cell activation - Langerhans, dendritic or macrophages - to drive T cell differentiation and B cell stimulation with increased Ig level;
- T helper cell mediated immune response with cytokine production to disturb proliferation and differentiation of epithelial cells and stimulate inflammatory reaction - Th2, Th1, Th17, Th22, Th9;
- innate lymphoid cell activation - ILC2 and ILC3 - to contribute to inflammatory response;
- granulocyte participation in pathogenesis - eosinophils and neutrophils - by their impact to tissue damage;
- sensory nerve activation to promote chronic pain and itch.

In more detail, the molecular mechanisms behind these phenotypes are described in Supplementary Material S1.

### 2.3 Intercellular layer: UC and AD maps

**The intercellular UC map**^2^ (Intercellular layer in Figure 1, Supplementary Material S3) is a three tissue-layer molecular network including 19 cell types. On the colonic epithelium layer, we show four cells, the most affected by the pathology: epithelial cells, goblet cells, primary afferent neurons, and enteroendocrine cells. The lamina propria layer, localised underneath the epithelium features Th naive cells, Th2, Th1, Th9, Th17, Th22, dendritic cells, ILC3s, macrophages, eosinophils, B cells, neutrophils and their production of mediators. Chemotaxis of innate and adaptive immune cells to lamina propria is shown in the blood vessel layer. The UC pathology is connected to the following factors: i) impaired epithelial barrier through affected colonic epithelial cells connected by tight junctions, ii) dysregulated immune response and iii) visceral pain^20^. Intercellular UC map demonstrates how epithelial barrier disruption induces immune dysregulation and other associated pathological mechanisms through cytokines, chemokines and other mediators activity. Detailed list of molecular events in the UC intercellular map with links to their location in the diagram online is described in Supplementary Material S3.

**The intercellular AD map**^3^ (Intercellular layer in Figure 1, Supplementary Material S2) represents a complex inflammatory network of 19 cell types and illustrates three tissue layers. The first epidermis layer represents cells involved the most in AD pathology. They are main skin epithelial cells: keratinocytes, Langerhans cells, inflammatory dendritic epidermal cells and pruriceptive primary afferent neurons. The second dermis layer contains different Th subsets such as Th naive cells and their differentiation into Th2, Th1, Th17 and Th22. Additionally, the diagram illustrates dermal dendritic cells, ILC2s, mast cells, eosinophils, B cells, neutrophils and others and their cytokine milieu. The third layer demonstrates chemotaxis of innate and adaptive immune cells from the peripheral blood to the skin. This molecular network of the AD map highlights how skin barrier dysfunction initiates a cascade of Th-mediated inflammation, with pruritus exacerbating this cycle. This is visualised through cell-to-cell communication in the intercellular layer. Detailed list of molecular events in the AD intercellular map with links to their location in the diagram online is described in Supplementary Material S2.

### 2.4 Pathway layer: shared pathology mechanisms

Various IMIDs share similar inflammatory pathways of common mechanisms in dysregulation of the innate and adaptive immune system^21^. In the UC and AD disease maps, we demonstrate these common mechanisms of UC and AD in the pathway layer in detailed pathways (Pathway layer in Figure 1, Supplementary Material S4). These pathways include NFkB, JAK-STAT, MEK-ERK signalling pathways in colonocytes (UC) and keratinocytes (AD), initiated by IL-13, IL-17A, IL-22, and IFNG, derived from immune cells. This is followed by downregulation of tight junction proteins and upregulation of alarmins, S100 proteins and IL-8. Another set of pathways involves IRAK-TRAF6-MAPKs activated by IL-33, as well as JAK-STAT6-GATA3 signalling activated by IL-4 in Th2 cells. This in turn upregulates Th2 cytokines such as IL-13 and IL-5. Finally, IRAK and NFkB signalling in Th1 cells initiated by IL-18 released from epithelial and dendritic cells and JAK-STAT1-TBX21 autocrine loop increases IFNG production.

A detailed list of mechanisms shown in submaps for various cell types can be found in Supplementary Material S4.

### 2.5 Exploration of the UC and AD maps

UC and AD maps are computational resources, supporting reproducible visualisation and analysis of omics data to interpret their contents in the context of disease. Here we introduce use cases of such applications, which can be reused in time when new datasets or map contents appear.

#### 2.5.1 Visual exploration of omics data

To show how gene variants might impact downstream molecular events represented in the maps, we gathered genes associated with UC from the Open Targets Genetics database^22^. These genes include those with variants, such as single nucleotide polymorphisms (SNPs), linked to UC. They were formatted into a data overlay for the UC map to visualise overlaps with represented disease mechanisms. OpenTargets data overlays visualise affected genes in the map, informing about most relevant pathways and cell types concerning disease genetics. Similar dataset was prepared for the AD map.

Additionally, we collected information from the ExpressionAtlas (https://www.ebi.ac.uk/gxa/) for keywords “colitis” and “dermatitis” in humans. Results were normalised and visualised in both maps for further insight into molecular mechanisms affected in both disorders.

These genetic and expression data overlays are openly accessible in UC and AD maps (the user guide is available online at https://disease-maps.io/ucadmap).

#### 2.5.2 Programmatic comparison of the UC and AD maps contents

To identify common and specific mechanisms across UC and AD, we performed a systematic computational analysis of their content, following the methodology in Gawron et al.^23^. To this end, we first compared stable HGNC identifiers (HUGO Gene Nomenclature Committee, www.genenames.org) shared between the maps. UC map and AD map feature respectively 230 and 270 unique HGNC identifiers, with 131 shared between the two maps. Figure 5 illustrates the most similar diagrams between the two maps, while Table 1 highlights ten most shared HGNC symbols, and which diagrams of the two maps they share.

**Figure 5.**
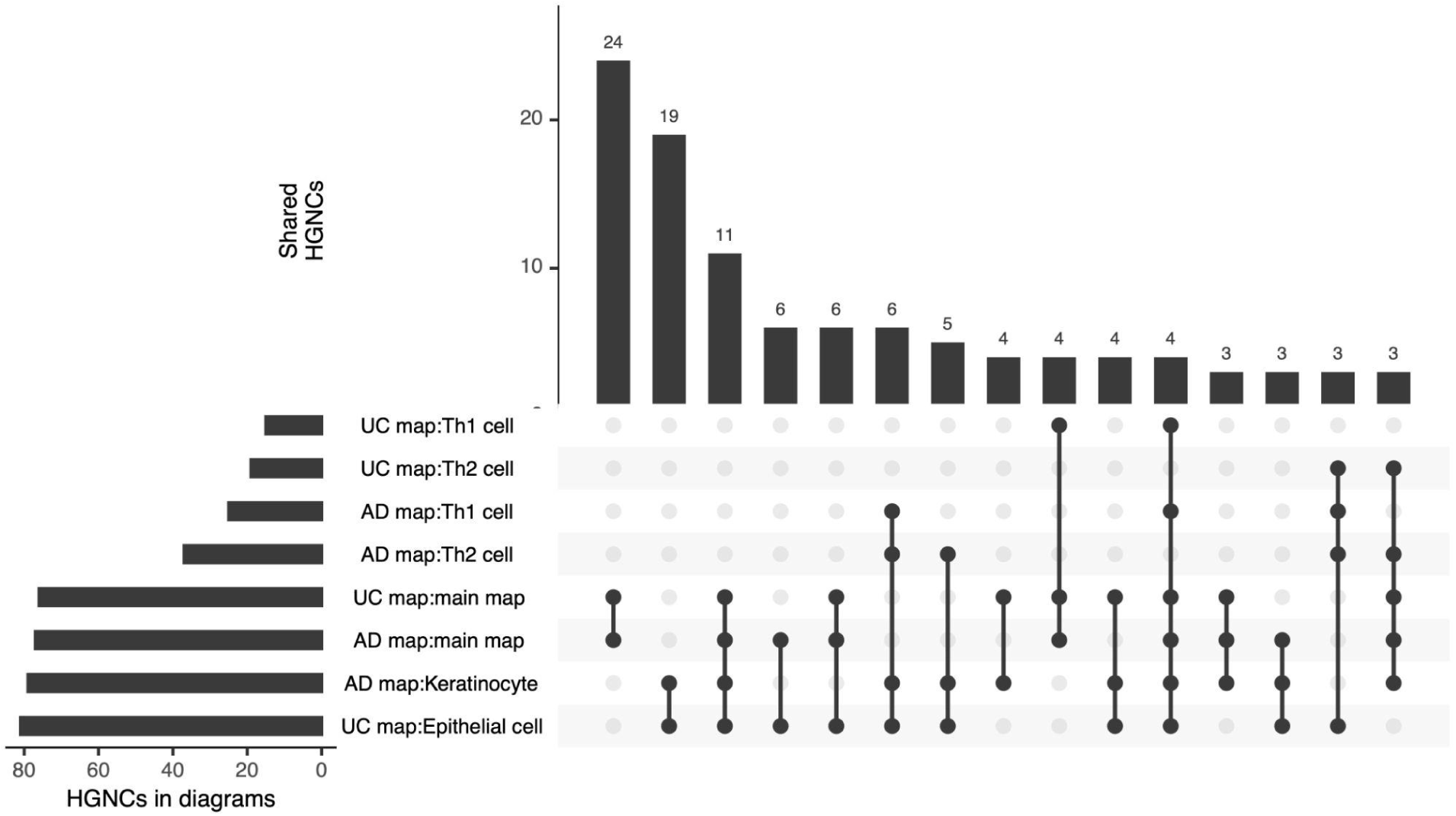
The number of shared HGNC identifiers between different diagrams of UC and AD maps. Similar diagrams with 131 common HGNC identifiers between the two maps are demonstrated. Data shown only for intersects of 3 and greater.

**Table 1.**
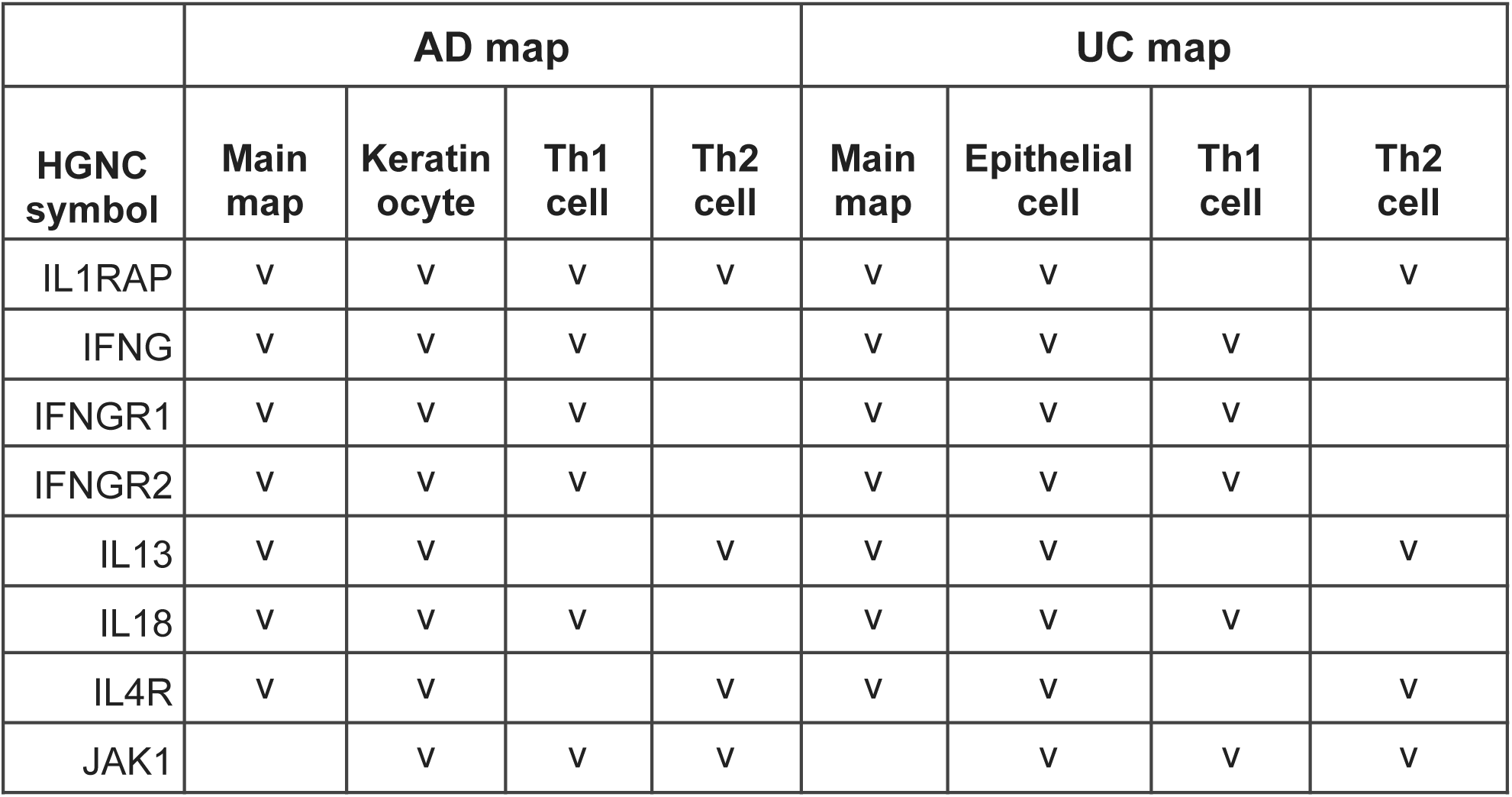
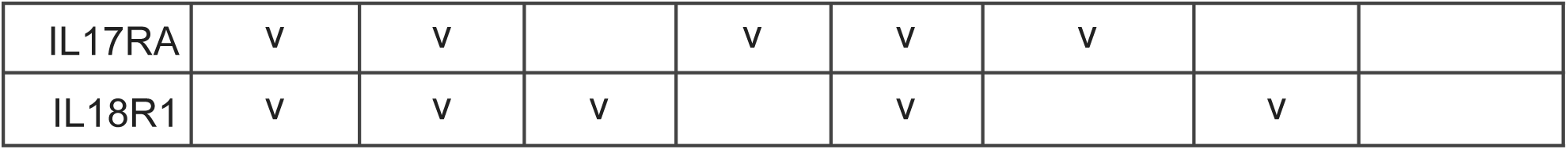
Gene symbols of top ten molecules common for UC and AD disease maps indicating in which diagrams they appear.

|  | AD map |  |  |  | UC map |  |  |  |
| --- | --- | --- | --- | --- | --- | --- | --- | --- |
| HGNC symbol | Main map | Keratinocyte | Th1 cell | Th2 cell | Main map | Epithelial cell | Th1 cell | Th2 cell |
| IL1RAP | v | v | v | v | v | v |  | v |
| IFNG | v | v | v |  | v | v | v |  |
| IFNGR1 | v | v | v |  | v | v | v |  |
| IFNGR2 | v | v | v |  | v | v | v |  |
| IL13 | v | v |  | v | v | v |  | v |
| IL18 | v | v | v |  | v | v | v |  |
| IL4R | v | v |  | v | v | v |  | v |
| JAK1 |  | v | v | v |  | v | v | v |
| IL17RA | V | V |  | V | V | V |  |  |
| IL18R1 | V | V | V |  | V |  | V |  |

To investigate similarities between cell types in both diseases, we retrieved lists of proteins associated with each cell type, and calculated Jaccard index values for these lists using HGNC symbols. Results of this comparison are illustrated in Figure 6. In the figure, cell type groups are indicated to highlight comparison between similar cell types. For the sake of clarity, values of 0.05 and lower were omitted in the figure. For instance, one can see high similarity between immune lymphoid cell types, and comparatively lower similarity between antigen presenting cells in both diseases. Gut epithelial cell in UC and keratinocyte in AD have similar molecular signatures, with overlap with profiles of Th2 and Th1.

**Figure 6.**
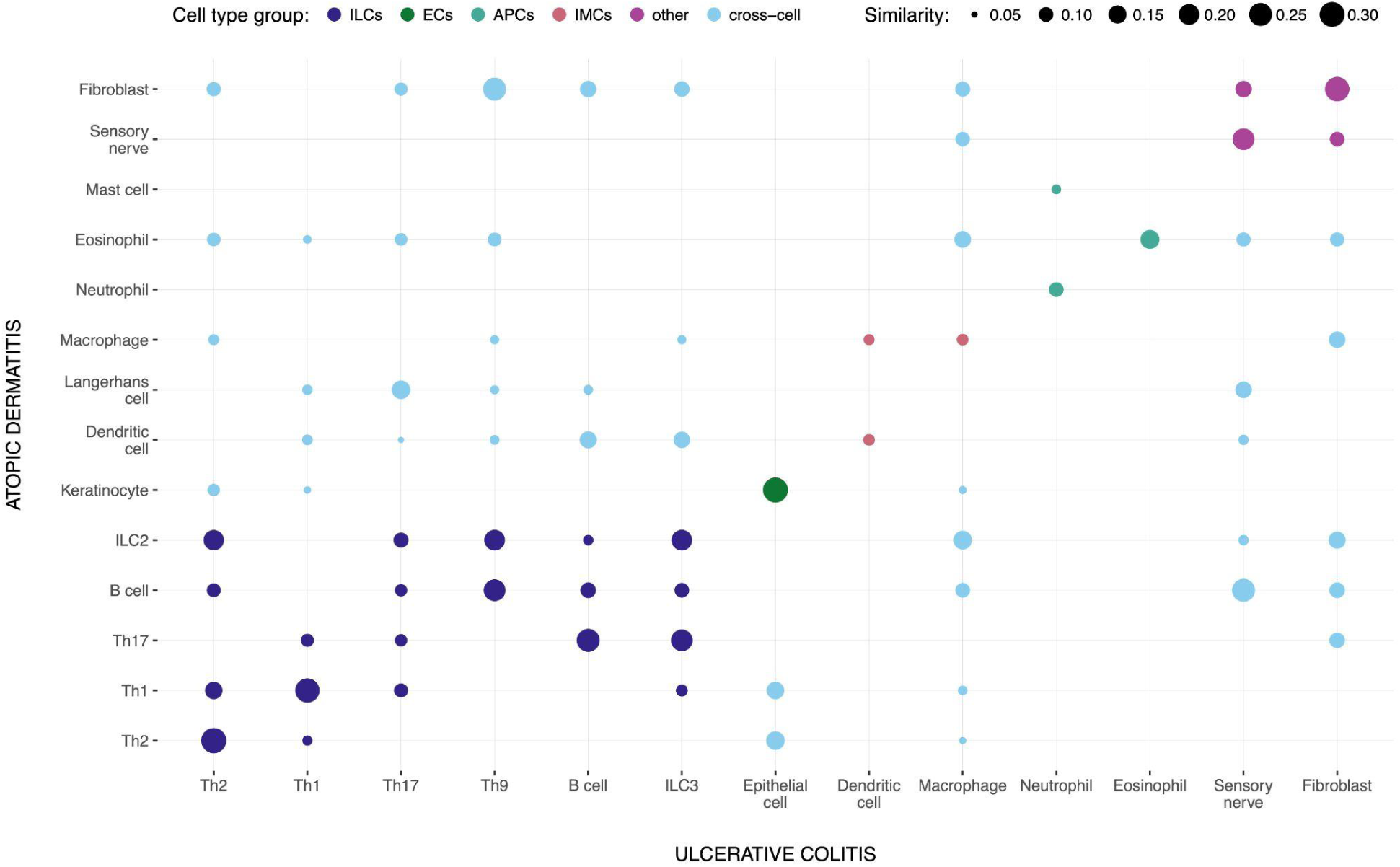
Comparison of molecular mechanisms in cell types in UC and AD based on corresponding disease maps. ‘Similarity’ denotes Jaccard index calculated pairwise for gene sets associated with respective cell types in UC and AD disease maps; values lower than 0.05 are omitted for clarity. Cells are grouped into categories. ‘Cross-cell’ comparisons indicate similarities between cells of different categories. ‘Cell type group’ abbreviations are: ILCs - immune lymphoid cells, ECs - epithelial cells, APCs - antigen-presenting cells, IMCs - immune myeloid cells.

To investigate the similarities between the structure of both maps, we calculated a score comparing interactions between the diagrams of both maps. Next, we grouped similar interactions and calculated overlap between these groups. This indicated 19 areas in both maps that have a significant level of similarity. Figure 7 illustrates two similar groups highlighted in both maps. The figure is offered for illustration purposes. The high-quality image with the ability for interactive zooming and further exploration is available online. Please see Supplementary Material S6.

**Figure 7.**
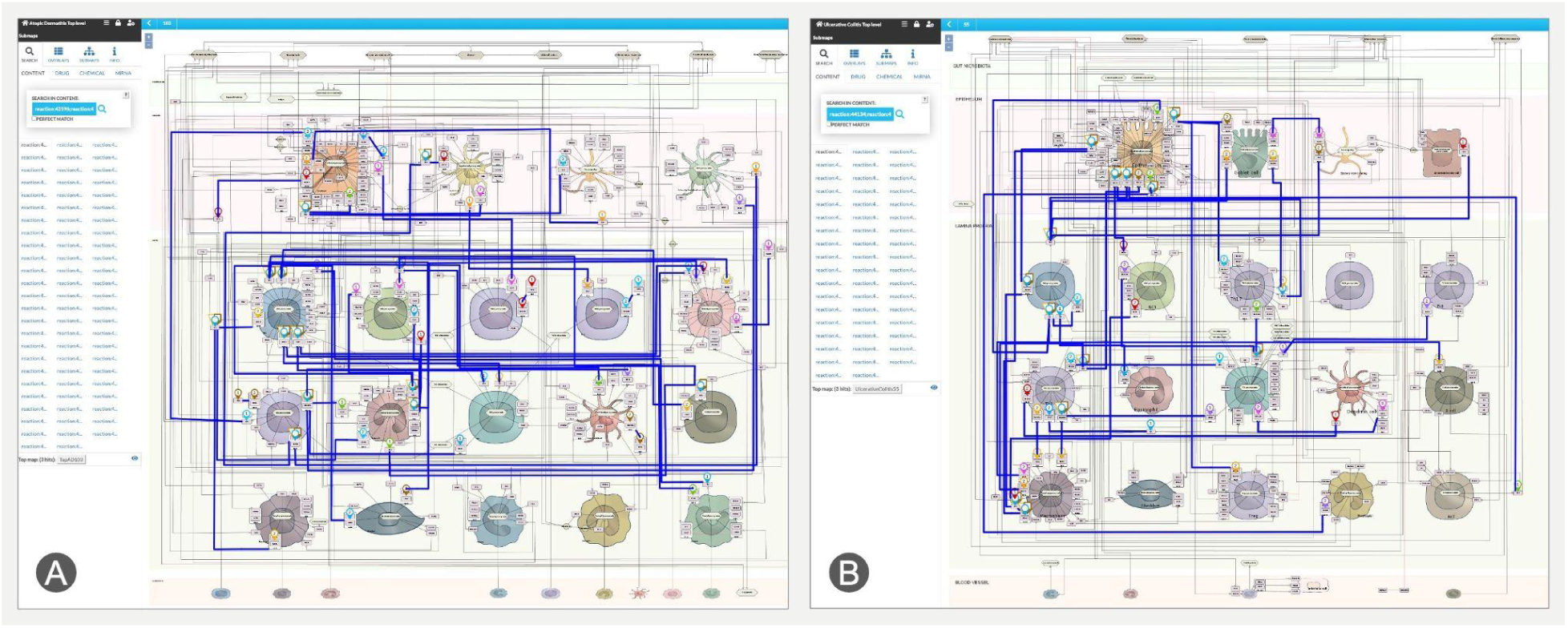
Visualisation of the overlapping pathology mechanisms of UC and AD in context of (A) AD and (B) in context of UC. This visualisation is available online for both disease maps along with other groups (see Supplementary material S6). Specific connections can be accessed from the table using the URLs provided.

Figure 8 illustrates a detailed example of such overlapping mechanisms for NFkB signalling cascade in Th1 cells in AD (left) vs epithelial cells in UC (right). This comparison reveals significant overlap in inflammation signalling pathways (impact on epithelial cells IL13, IL17, IL22 and IFNG, Th activation by IL6, eosinophil activation by IL5, Th1 differentiation with stimulation by IL12 etc.).

**Figure 8.**
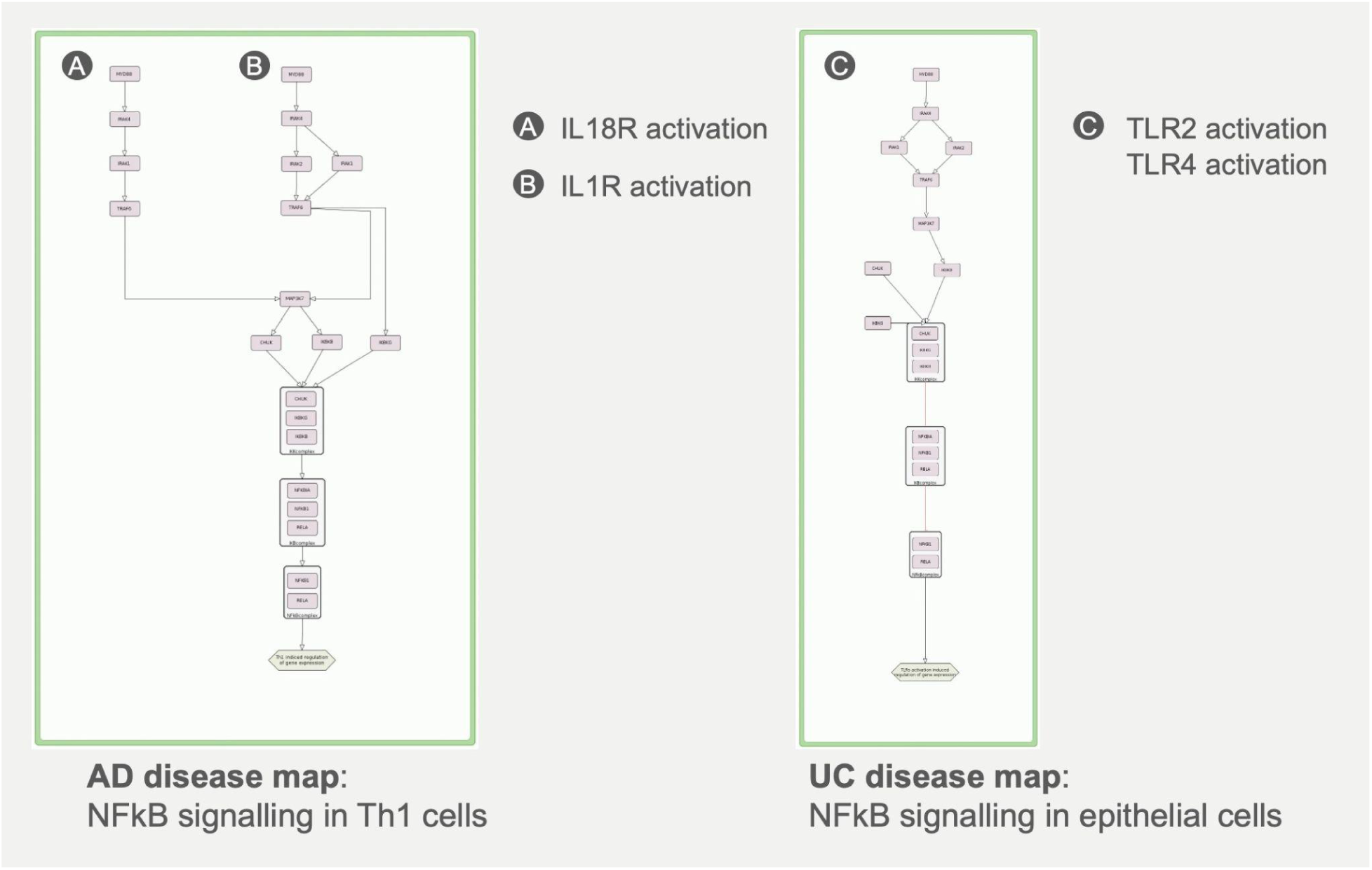
Comparison of NFkB signalling in AD and UC maps across different cell types shows different regulation of similar core mechanisms.

Overall, 19 groups of similar interactions include 411 of 825 interactions of the UC map, and 530 of 1238 interactions of the AD map, with TRAF6, MAP3K7, CHUK, IKBKB and IKBKG present across 7 or more groups. Please see Supplementary material S5 for a complete list of similar groups, their content and direct links in corresponding disease maps.

## 3. DISCUSSION

A systematic review of the literature on UC and AD reveals highly complex mechanisms underlying these disorders. By integrating this knowledge into a structured, visual, and computational resource, disease maps provide a powerful tool for advancing our understanding of these pathologies and enable their systematic comparison. This approach not only allows to tackle the complexity of available information but also facilitates hypothesis generation and potentially applications for identifying drug targets and therapeutic strategies.

### 3.1 Shared UC and AD mechanisms

UC and AD are both inflammatory disorders of epithelial surfaces, where UC affects colonic epithelium and AD - epithelial tissue of the skin. The pathology changes the structure and function of colonic epithelial cells^24^ and keratinocytes^25^ and may result in hyperplasia or other abnormal conditions - epithelial abnormalities in UC^9^ and barrier dysfunction in AD^8^. Both colonocytes and keratinocytes are influenced by cytokines IL-13, IL-4, IL-17, IL-22, IFNG and others, and the response to this influence is the expression of alarmins like IL-33^26,27^. IL-25 and TSLP are also important in AD^28,29^, while IL-8 for UC^30^. Both maps illustrate these mechanisms^45^ such in AD proinflammatory IL-13, derived from Th2, ILCs, mast cells and NKT cells that acts on keratinocyte through mTOR-AKT signalling, and downregulates FLG, essential for skin barrier^31^. Together with IL-4, it also downregulates Loricrin, Involucrin, FLG2, involved in formation and maintenance of epidermis^32,33^. Similarly, in UC, IL-13 downregulates tight junction proteins Claudin8, Occludin^34^, and tricellulin^35^, critical for epithelial barrier formation, and upregulates Claudin2^36^, a pore-forming protein responsible for increased permeability for small cations. This way, in both pathologies IL-13 is responsible for a loss of the epithelial barrier integrity.

Dysregulation of T helper cell subsets contributes to the pathogenesis of UC and AD. Acute AD is characterised by Th2 domination with IL-4, IL-13 and IL-5 release^37^. However chronic AD is characterised by Th1 activation, as well as Th17 and Th22 subsets^38^. Asian, paediatric, and intrinsic endotypes of AD also have a strong association with Th17 and production of IL-17^39^. In turn, UC is associated with the response of Th1 subset, producing IFNG and TNF, and Th17 subset, producing IL-17, that mainly promote inflammation and tissue damage in the gut^40^. However some endotypes of UC show activation of Th2 subset with IL-4, IL-13 and IL-5 production as well^36^. Th9 subset is involved in UC pathology with impact on the epithelial cell^41^. Both maps demonstrate^45^ how IL-4 signalling in Th2 cells stimulates increased inflammatory response through JAK STAT6^42^, GATA3^43^ in UC and in AD^44,45^. IL-18, released from keratinocytes^46^ and inflammatory dendritic epidermal cells^47^, mediates Th1 response to enhance IFNG production through IRAK and NFkB in AD^48,49^. Similarly, in UC, IL-18 derived from colonocytes^50^ or dendritic cells^51^, mediates inflammatory response acting on Th1 with increased IFNG production^52^. IFNG itself increases IFNG Th1 expression acting in an autocrine loop through JAK-STAT1-TBX21 as in AD^53^ and in UC^54,55^.

Among the immune cell subsets, pathology of UC and AD shows innate lymphoid cell activity. While different subsets of ILC play their roles in the pathology of two diseases, activation of type 2 innate lymphoid cells (ILC2) and their cytokine production (IL-4, IL-13 and IL-5) contribute to the chronic inflammation and tissue damage in AD^56^, and type 3 innate lymphoid cells (ILC3) with IL-17 release play a significant role in the pathogenesis of UC^57^.

Other cell types involved in both disorders are granulocytes. Eosinophils are particularly prominent in AD, where they contribute to tissue inflammation and damage through the release of IL-12, exacerbating the skin inflammatory response^58^. In UC, neutrophils release neutrophil extracellular traps (NETs) that can lead to tissue injury and chronic inflammation^59^.

Finally, sensory nerve activation plays a crucial role in the development of chronic pain associated with both conditions. In AD, various mediators, IL-31 and IL-13 among them, can sensitise sensory nerves in the skin, resulting in heightened pain perception and itch^60^. In UC, inflammatory processes in the gut can activate sensory neurons, contributing to visceral pain by IL-13 derived from Th2 and NKT cells^61^. Pathology mechanisms of UC and AD shown in detail in the intercellular and pathways layers, are described in Supplementary materials S2, S3, S4.

### 3.2 Computational analysis

UC and AD disease maps are computational resources, and computational analysis of their overlap (Figure 6, Table 1) reveals shared biomarkers and common pathological pathways. Table 1 summarised the most common proteins for both diseases, supporting the inflammation-related signature. These proteins include key Th2 and Th1 cytokines and their receptors IL13 and IL4R, IFNG and IFNGR1/IFNGR2. This reinforces our earlier discussion on the involvement of both Th2 and Th1 immune responses in the pathogenesis of UC and AD. Both IL13 and IL4 are potential plasma biomarkers for AD^62^. In UC, serum IL13 is used as a potential biomarker^63^. The list includes IL18, associated with the disease severity. It promotes the differentiation of Th0 into Th1 cells and stimulates Th1 to produce IFNG and is associated with severity in both AD^64^ and UC^65,66^.

Figure 6 illustrates similarities between cell types in both diseases. As expected, immune lymphoid cells show high similarity for Th2, Th1, B cells and ILCs between diseases. As for cells from different categories, ILC2 in AD and Th2 in UC have close molecular signatures of proinflammatory cytokines. ILC2s in AD, induced by alarmins, express IL-4, IL-13 and IL-5^56,67,68^. Th2s are activated in some endotypes of UC with IL-4, IL-13 and IL-5 production as well^36^. In both diseases these molecules impact on epithelial cell damage and eosinophil activation^36,56,69,70^. Another interesting pair of cells from different groups with similar molecular patterns is Th17 in AD and ILC3 in UC. ILC3s in UC with IL-17 release contribute to intestinal inflammation^57^. Some AD endotypes also show IL-17 production by Th17s^39^ which increases tissue damage and provokes chronic skin inflammation^71^. Immune myeloid cells - eosinophils, neutrophils - show similar activity for both diseases, as well as fibroblasts and sensory nerve endings involvement that support our previous observations about mechanisms of tissue injury, chronic inflammation and pain^58–61^.

Finally, computational analysis indicated specific areas in both maps with similar networks of molecular mechanisms. Figure 8 shows one of 19 groups of such interactions, featuring similarities of NFkB signalling in UC and AD maps across different cell types. It indicates different regulation of similar core mechanisms in UC and AD. Despite various signals triggering NFkB activation, we identify similar events following activation of IL1R in Th1 cells in AD and Toll-like receptors in epithelial cells in UC. Our analysis points to upregulation of TRAF6, IKBKG and MAP3K7 associated with response to IL1R ligands (IL18, IL1B) and TLR ligands (LPS) but not to other triggers, as supported by^72–74^.

Although significant efforts have been made to identify therapeutic agents for both UC and AD, only a limited number of successful cases have been reported. Notably, treatment with tofacitinib^75^ and upadacitinib^76^ in patients with coexisting UC and AD has demonstrated clinical efficacy. Both drugs are JAK inhibitors that target the JAK-STAT intracellular signaling pathways which play a crucial role in the pathogenesis of both conditions. However, the management of UC and AD remains complex, highlighting the need for novel, mechanism-based therapeutic strategies.

Computational analysis of disease maps identifies disease-specific and shared biomarkers for UC and AD which gives us a better picture of their association and new possible targets for treatment options. Although the two diseases are known to be related before, our work demonstrates clearly their overlapping mechanisms and common pathways across different cell types. Workflows used to compute these comparisons can be repeated, which will produce new insights when both resources will be further developed by the research community. This will aid ongoing investigations into UC and AD comorbidities, facilitate data interpretation and advance research towards addressing common and specific mechanisms of these disorders for novel treatment strategies. Moreover, with other disease maps and pathway databases being available, similar studies in adjacent areas are feasible as well.

### 3.3 Perspectives and future updates

The UC and AD disease maps were mainly developed through a systematic curation of articles available in PubMed to show our current knowledge about the disease mechanisms. Since new UC- and AD-related articles are continuously published, we plan to regularly check and update the diagrams to ensure that they remain relevant. The structure of the resource allows performing such updates, and the map building effort is supported by the ImmUniverse consortium (https://www.immuniverse.eu/).

The content of the maps was evaluated by domain experts and the resource was extended following feedback provided by the ImmUniverse consortium. While we recognise that it is challenging to comprehensively collect all published information on a topic, the systematic methodology^19^ allows to cover major disease areas and the mechanisms behind them.

## 4. MATERIALS AND METHODS

### 4.1 Map construction

Information on the molecular mechanisms of UC and AD was extracted from biomedical literature and encoded into diagrams (disease maps). CellDesigner (https://www.celldesigner.org) is a diagram editor that we used to create UC and AD disease maps. It follows Systems Biology Graphical Notation (SBGN) standard^77^. This editor is connected to the MINERVA visualisation platform^78^ with access for visualisation and exploration.

#### 4.1.1 Literature search

Our curation process is described in the recent community guideline publication^19^. We collect evidence from biomedical publications which confirm the relevance of a molecule interaction for UC or AD diseases. As evidence for each interaction we use experimentally validated data from disease-relevant papers and pathway databases. Evidence is integrated from PubMed and PubMed Central searches, from pathway databases such as Reactome^79^ and KEGG^80^, and annotation databases such as UniProt (https://www.uniprot.org) and CHEBI (https://www.ebi.ac.uk/chebi). We provide confirmation for each molecular interaction and give references to the corresponding publications. All interactions are manually curated, and evidence is provided in the form of publication identifiers - PMID or DOI. Eight hundred and three papers were selected for this project, being 325 and 478 for, respectively, UC and AD intercellular maps and submaps. Firstly, we reviewed and analysed key papers which were suggested by domain experts from the ImmUniverse consortium. Then a broader search was performed to integrate not only disease-specific information from publications but also more general relevant experimental data for immune inflammatory conditions to fill missing elements of the disease pathology. When possible, we use pathway databases to fill gaps in downstream pathway events.

We discussed the scope of the maps with domain experts. We updated the maps following feedback provided by the ImmUniverse consortium to extend the maps with genes found to be linked to UC and AD. To integrate suggested genes into the maps, we follow our protocol which includes a confirmation of every step of the disease mechanism with definite experimentally validated data for a specific gene or protein, specific cell type and tissue in disease-relevant papers.

#### 4.1.2 Encoding interactions into diagrams

Information about molecule interactions from publications is manually integrated, reviewed and represented in a diagrammatic form^81^. To create, edit or update UC and AD disease maps, the CellDesigner diagram editor is used for developing disease maps^82^. It supports extensive diagrams, follows System Biology Graphical Notation (SBGN) logic and makes it possible to draw SBGN-compatible diagrams in Process Description and Activity Flow type of languages. As a graphical standard, we use SBGN^77^ ActivityFlow language to show molecules and communications between them.

#### 4.1.3 Annotation of map elements and interactions

Each entity on the maps is identified and has a link to the corresponding external database according to the object type. We identify entity type (protein, simple molecule, etc.) and use a standard name for every entity or compartment in CellDesigner. All proteins, RNAs and genes are named according to the official symbols of HUGO Gene Nomenclature Committee (HGNC) and annotated with the corresponding UniProt IDs. Simple molecules are identified via Chemical Entities of Biological Interest database (CHEBI). Phenotypes are identified by GO biological process terms and the corresponding IDs (Gene Ontology) in case they represent biological processes, or Medical Subject Headings (MeSH) in case they represent disease-related terms. Automatic annotation in MINERVA is possible for proteins, RNAs and genes as soon as they have HGNC names. Manual annotation is needed in cases where generic names are used or there are many synonyms, so entities are not recognised automatically by MINERVA, in particular for metabolic entities and phenotypes. All interactions are annotated with PMIDs or DOIs. Disease-specific evidence is drawn from the literature, and annotating map interactions requires maintaining a substantial repository of articles. All articles used for this project are stored in a Zotero library.

### 4.2 Maps availability and visualisation in the MINERVA platform

To represent the molecule interaction network, we use the MINERVA (Molecular Interaction NEtwoRks VisuAlization) platform^78^, a web service for disease map content exploration and data interpretation. The UC and AD comparison diagram is accessible at https://imi-immuniverse.elixir-luxembourg.org. Users can browse the resource by clicking on the content to see maps description, search for map elements (molecules, connections, publications), or explore data by overlaying datasets (transcriptomics, gene variant data, etc.) over UC and AD disease maps, which will be transformed into a colour representation on the maps.

### 4.3 Construction of data overlays

Data overlays for UC and AD maps were constructed using Open Targets^83^ and Expression Atlas^84^ resources. Open Targets platform was queried for genetic variants associated with UC (EFO_0000729) and AD (EFO:0000274) using their API on 27th of September 2024. These lists were compiled into two overlays indicating genes affected by at least one variant.

Expression Atlas was queried for “colitis” and “dermatitis” condition terms for “homo sapiens” set as species on 28th of May 2024. In both datasets, log fold change was scaled to [-1;1] range for visualisation in the MINERVA Platform.

### 4.4 Computational analysis of overlaps between UC and AD maps

UC and AD maps were compared following the workflow introduced in^23^. HGNC symbols were used as element identifiers. Pairwise similarity between interactions of both maps were calculated using the same similarity metric as in the original article. Only interactions having similarity of at least 0.7, and at least two similar elements were retained. For these interactions in both maps, strongly connected components (SCCs) were identified. SCCs were then compared pairwise between the maps, and pairs with 5 or more similar reactions were kept as groups of similar interactions.

## CONCLUSION

UC and AD were known to be related, with AD patients exhibiting an increased risk of developing UC. Our study focuses on shared and unique molecular mechanisms of the diseases. We compare them by constructing UC and AD disease maps followed by a computational analysis. We show that despite different manifestations, both conditions exhibit common epithelial barrier dysfunction, immune dysregulation (such as Th2, Th1 and ILC response) and sensory nerve activation. Key inflammatory pathways were identified in both diseases such as NFκB, MEK-ERK, IRAK-TRAF6-MAPK, JAK-STAT6-GATA3 and JAK-STAT1-TBX21, showing overlapping mechanisms of chronic inflammation and tissue damage. Moreover, our computational analysis identified shared biomarkers such as IL-13, IL-4R, IFNG, and IL-18. The presence of common signaling cascades and biomarkers supports the potential for drug repurposing where treatments effective in one disease may offer therapeutic benefits in the other.

Disease maps presented here are community-constructed knowledge repositories that offer translation of visually legible and pathology-specific pathway diagrams into analysis and modelling resources. The workflows can be reused with further results when applied to new projects and when the map is updated based on new fundings.

By providing an interactive and open-access resource, the UC and AD disease maps can support future research, personalised medicine, and predictive modeling for IMIDs. As new findings emerge, continuous updates to these maps will further refine our understanding of disease mechanisms, aiding in the development of novel treatment strategies and improved patient outcomes.

## DISCLAIMER

Any dissemination of results reflects only the author’s view and the JU is not responsible for any use that may be made of the information it contains.

## ACKNOWLEDGMENTS

The work presented in this paper was supported by the ELIXIR Luxembourg tools and services.

## Footnotes

1 https://imi-immuniverse.elixir-luxembourg.org

2 https://imi-immuniverse.elixir-luxembourg.org/minerva/index.html?id=UCmaps31-01-25

3 https://imi-immuniverse.elixir-luxembourg.org/minerva/index.html?id=ADmaps31-01-25

4 https://imi-immuniverse.elixir-luxembourg.org/minerva/index.html?id=UCmaps31-01-25

5 https://imi-immuniverse.elixir-luxembourg.org/minerva/index.html?id=ADmaps31-01-25

