## Supplementary Materials for "Association of ulcerative colitis with atopic dermatitis: identification of shared and unique mechanisms by construction and computational analysis of disease maps"

### Supplementary Material S1

Following mechanisms, shown in the comparison diagram (Figure 2) define the connecting and distinguishing characteristics of AD and UC.

**Epithelial cells involvement** (Figure 2): AD and UC are both inflammatory disorders of epithelial surfaces, where AD affects the epithelial tissue of the skin and UC - colonic epithelium. As a protective layer, skin epidermis provides a barrier to various antigens and triggers [19043850]. Skin epithelial cells keratinocytes are the most abundant cell types of human epidermis, and present 90% of epidermis cells. They form the epidermal barrier by their tight junctions [11889141]. Another example of such a barrier is intestinal epithelium. Colonic epithelial cells colonocytes play a central role in maintaining colon homeostasis and its imbalance [30498100]. Colonocytes also form tight junctions to perform a barrier function [35621376]. The pathology changes the structure and function of keratinocytes [28583445] and colonic epithelial cells [11729111] and may result in hyperplasia or other abnormal conditions. Barrier dysfunction is one of the initial steps in the development of atopic dermatitis [34639001], and keratinocytes play a critical role in its pathology mechanisms. Epithelial abnormalities present in ulcerative colitis patients and underlie the development of mucosal inflammation and its chronicity [31278430]. Thus, keratinocytes and colonic epithelial cells are the most affected cells in immune-inflammatory disorders, and mechanical processes associated with them are very important. Both of these cells are influenced by cytokines IL-13, IL-4, IL-17, IL-22, IFNG and others, and the response to this influence is the expression of alarmins like IL-33 [33011245, 19913053]. IL-25 and TSLP are also important in AD [32179159, 38198924], while IL-8 for UC [12474223]. More detailed mechanisms are described in corresponding submaps. Antigens are directly connected to epithelial and antigen-presenting cells to trigger inflammatory processes.

**Specific types of T lymphocytes and their cytokines** (Figure 2): Dysregulation of T helper cell subsets contributes to the pathogenesis of AD and UC. Acute AD is characterised by Th2 domination with IL-4, IL-13 and IL-5 release [31182684]. However chronic AD is characterised by Th1 activation, as well as Th17 and Th22 subsets [30685456]. Asian, paediatric, and intrinsic endotypes of AD also have a strong association with Th17 and production of IL-17 [31465744]. In turn, UC is associated with the response of Th1 subset, producing IFNG and TNF, and

Th17 subset, producing IL-17, that mainly promote inflammation and tissue damage in the gut [23446770]. However some endotypes of UC show activation of Th2 subset with IL-4, IL-13 and IL-5 production as well [16083712]. One of the differences between these two conditions is the involvement of the Th9 subset in UC pathology, IL-9 produced by them and its impact on the epithelial cell [24908389].

**Innate lymphoid cell types** (Figure 2): Among the immune cell subsets, pathology of AD and UC shows innate lymphoid cell activity. There are different subsets of ILC that play their roles in the pathology of two diseases. Activation of type 2 innate lymphoid cells (ILC2) and their cytokine production (IL-4, IL-13 and IL-5) contribute to the chronic inflammation and tissue damage in AD [32179159]. Type 3 innate lymphoid cells (ILC3) with IL-17 release play a role in the pathogenesis of UC [30962426].

Various **granulocytes**, including eosinophils and neutrophils (Figure 2), play critical roles in AD and UC. Eosinophils are particularly prominent in AD, where they contribute to tissue inflammation and damage through the release of IL-12, exacerbating the skin's inflammatory response. In UC, neutrophils release neutrophil extracellular traps (NETs) that can lead to tissue injury and chronic inflammation.

**Sensory nerve** activation (Figure 2) plays a crucial role in the development of chronic pain associated with both conditions. In AD, various mediators, IL-31 and IL-13 among them, can sensitise sensory nerves in the skin, resulting in heightened pain perception and itch [24373353]. In UC, inflammatory processes in the gut can activate sensory neurons, contributing to visceral pain [36749569].

### Supplementary Material S2

#### Description of the AD intercellular map

Pathology mechanisms of AD are driven by three interrelated factors: skin barrier disturbance, Th response, and pruritus [31902917]. Consequently, main molecular events in AD include activation of epidermal keratinocytes, immune cells, and peripheral sensory nerves.

**Skin barrier disruption** is the result of epidermal hypersensitivity to external triggers, which permits external pathogens and allergens to penetrate through the skin barrier and activate keratinocytes, leading to the production of alarmins

TSLP, IL-25, and IL-33 [12055625, 32179159]. These signals mediate intercellular mechanisms between the epidermal keratinocytes and immune cells. In response, various cytokines and chemokines act on keratinocytes and provoke epidermal barrier dysfunction. IL-4 and IL-13 are directly involved in reducing filaggrin expression in the skin [33865911]. It results in penetration of antigens and allergens to maintain inflammation as filaggrin is a protein that maintains barrier integrity. IL-13 and IL-4 trigger keratinocytes to release various cytokines and chemokines, leading to altered inflammatory processes and inhibited barrier function. These cytokines are able to reduce the expression of barrier proteins such as loricrin or involucrin making the epidermis more disturbed by antigens [18166499]. Changes in filaggrin expression and tight junctions provoke barrier disruption, which leads to abnormal permeability of the skin [16550169, 35628125]. IL-33 acts on keratinocytes in an autocrine manner and negatively regulates the expressions of filaggrin and other skin barrier proteins [29534857]. A high level of IL-22, produced by Th22, downregulates filaggrin expression as well, reduces keratinocyte differentiation and induces epidermal hyperplasia [21564072]. Hyperproliferation of keratinocytes can also be triggered by dermal fibroblasts in AD that contribute to barrier dysfunction [22684102].

**Immune response** is facilitated essentially by immune cells that detect and process pathogens. These antigen-presenting cells - Langerhans, IDEC and dendritic cells, as part of the innate immune system, take up antigens penetrating through the skin barrier and stimulate Th cell differentiation [15131579]. Cutaneous dendritic cells are activated by TSLP and by external protein antigens encountering the skin [12055625]. These cells express MHC class II and costimulatory molecules, promoting the differentiation of naive T cells into Th2 cells and other Th cells and the attraction of them into the skin [22548207]. TSLP primes Langerhans cells in the epidermis [17320941]. Activated macrophages also act as antigen-presenting cells and contribute to angiogenesis and lymphangiogenesis in AD pathology [22574929]. IL-33, produced by keratinocytes, is shown to promote proliferation, activation, and recruitment of Th2 cells, activate mast cells, ILC2 and basophils [22277940]. Th2 cells produce proinflammatory cytokines such as IL-4 and IL-13 - to reduce filaggrin in keratinocytes and disturb barrier function, IL-31 - to act on sensory nerve endings and stimulate scratching, and IL-5 - to activate eosinophils [11754819, 24373353, 14600427]. IL-4 also controls the differentiation of Th2 and increases B cell IgE production [7918008, 17267416, 22430776]. IL-13 and IL-4 are produced not only by Th2 cells, but also by ILC2 and mast cells [12145428, 7918008,

32179159]. ILC2 can be induced by alarmins directly to express them [32179159, 24323357, 23363980]. Basophils function as a source of IL-4 as well [33662369]. IL-5 expressed by Th2 cells activates eosinophils and attracts them to the skin via CCL26 [16045735]. ILC2 is another source of IL-5 in AD [23603794]. Appearance of Th1, Th22 and Th17 subsets shows a transition from the acute to the chronic stage with expression of IFNG in patients with chronic lesions [28266782]. Their cytokine production (IL-17, IL-22, IL-26) also contributes to the pathogenesis of AD [27588419, 19439349, 31465744].

**Pruritus** is one of the main symptoms in patients with AD [30471983]. Epidermal keratinocytes can be activated by scratching, infection or irritants in response to tissue damage because of pruritus [32738956]. Various pruritogens induce itching and scratching: IL-4, IL-31, IL-13, TSLP [28890086, 24094650, 24373353, 34391816] as well as histamine produced by mast cells [35113811]. Itch is mediated by pruriceptive sensory nerve endings, which transmit the pruritic signal to the brain and result in pain [28890086]. Scratching leads to further tissue damage and activation of keratinocytes, making an inflammatory loop in AD patients [30446754]. Neutrophils are involved in chronic pruritus [31631836].

### Supplementary Material S3

#### Description of the UC intercellular map

The UC pathology is connected to a few major factors: impaired epithelial barrier through affected colonic epithelial cells connected by tight junctions, dysregulated immune response and visceral pain [27914657]. The second factor involves innate and adaptive immune cells infiltrating the lamina propria. In our disease map we show numerous changes in molecular mechanisms, among which the most significant are the following.

**Mucosal and epithelial barrier defects**, mostly impacting colonocytes, contribute to the inflammatory cascades. Activated Toll-like receptors TLR2 and TLR4 in colonocytes are responsible for the disruption of epithelial tight junctions and increased permeability of the barrier [12538701, 10623846]. Epithelial cells produce large amounts of cytokines and chemokines that participate in pathology mechanisms of tissue injury in UC. Goblet cells produce and secrete protective mucins to prevent intestinal inflammation caused by penetrating pathogenic bacteria. A decrease in the number of goblet cells and inhibited mucin secretion is reported in UC patients [30914450]. Interleukins IL-13 and IL-4, produced by

Th2 or NKT cells, induce apoptosis of epithelial cells [12433369]. IL-13 also acts on nerve endings and causes hypersensitivity in UC patients [36749569]. Cytokines TNF and IFNG produced by various cells contribute to cytotoxicity and have direct effects on epithelial barrier integrity [17923086, 21078084]. IL-9 expression by Th9 impairs colonocyte proliferation and the intestinal barrier work [24908389]. IL-10, IL-22 and IL-2 can be antiinflammatory factors in UC [4367025, 23715173]. They enhance the intestinal epithelial barrier integrity and tissue repair. IL-22 protective mechanism works through stimulation of mucus production by goblet cells [18172556]. IL-18 on one hand protects against UC in the acute stage of mucosal immune response [31348891]. On the other hand, IL-18 decreases goblet cell development and their mucus production [26638073].

**Immune response** maintenance is the result of impairments of tight junctions in the mucosal layer and increased uptake of luminal antigens. Dendritic cells are first responders and have enhanced expression of costimulatory molecules [22619032]. Activated dendritic and other antigen-presenting cells initiate the differentiation of naive T cells into Th2 cells, Th1, Th17, Th2 or Th9. Th2 produces proinflammatory cytokines IL-5, IL-4, and IL-13 [16083712, 30383225, 8757634]. IL-13, produced by NKT cells, also takes part in epithelial injury [15146247]. Th cells also differentiate into Th1, which expresses IFNG. It has been shown that neutralisation of IFNG completely annuls colitis [26883725]. TNF and IL-21 production by Th1 is significant for this pathology as well [20677335, 30946450]. Macrophages are also reported to be involved in the TNF-mediated UC [21875498]. Th17 and Treg differentiation is regulated by IL-18 [31348891] and IL-2 [30944470], which suppress inflammation in UC. Th9 cells, which are induced by IL-36 derived from the epithelium, express IL-9 and decrease cellular proliferation and weakens the intestinal barrier function [24908389]. Innate lymphoid cells ILC3s contribute to chronic intestinal inflammation and show increased cytokine production of IL-17 and IL-22 as a result of IL-23-driven inflammatory response [32332067]. Activated neutrophils accumulate in the colonic tissue in UC. Stimulated by TNF they form neutrophil extracellular traps as a part of UC-related inflammation [30715224]. Increased IgG level expressed by B cells is reported in UC patients. This causes IgG-dependent inflammation [27914657].

**Chronic pain** is a common symptom in UC [22298998]. Colonic sensory neurons are sensitised by IL-13 derived from Th2 and NKT cells [36749569]. One of key mediators of neuronal excitability in UC is TNF [20427396]. Its production is elevated in UC in various immune cells and TNF receptors are expressed by neurons innervating the colon. Also serotonin signalling via HTR3 in mucosal nociceptors contribute to the visceral hypersensitivity in UC [22330338].

**3.2** Shared mechanisms, indicated in the comparison level, are demonstrated in more detail in the intercellular layer of the AD and UC disease map (Figure …)

Pathology mechanisms of AD are driven by three interrelated factors: skin barrier disturbance, Th response, and pruritus [31902917]. Consequently, main molecular events in intercellular AD map include activation of epidermal keratinocytes, immune cells, and peripheral sensory nerves.

Similarly, in the UC pathology inflammatory cascades that can be initiated by mucosal and epithelial barrier defects and environmental factors, leads to chronic inflammation and pain [27914657]. Intercellular UC map shows numerous changes in those molecular mechanisms.

**Skin barrier disruption in AD** is the result of epidermal hypersensitivity to external triggers, which permits external pathogens and allergens to penetrate through the skin barrier and activate keratinocytes, leading to the production of alarmins TSLP, IL-25, and IL-33 [12055625, 32179159]. These signals mediate intercellular mechanisms between the epidermal keratinocytes and immune cells. In response, various cytokines and chemokines act on keratinocytes and provoke epidermal barrier dysfunction. IL-4 and IL-13 are directly involved in reducing filaggrin expression in the skin [33865911]. IL-13 and IL-4 trigger keratinocytes to release various cytokines and chemokines, leading to altered inflammatory processes and inhibited barrier function [18166499]. Changes in filaggrin expression and tight junctions provoke barrier disruption, which leads to abnormal permeability of the skin [16550169, 35628125].

**Mucosal and epithelial barrier defects in UC**, mostly impacting colonocytes, contribute to the inflammatory cascades. Activated Toll-like receptors TLR2 and TLR4 in colonocytes are responsible for the disruption of epithelial tight junctions and increased permeability of the barrier [12538701, 10623846]. Epithelial cells produce large amounts of cytokines and chemokines that participate in pathology mechanisms of tissue injury in UC. A decrease in the number of goblet cells and inhibited mucin secretion is reported in UC patients [30914450]. Interleukins IL-13 and IL-4, produced by Th2 or NKT cells, induce apoptosis of epithelial cells

[12433369]. Cytokines TNF and IFNG produced by various cells contribute to cytotoxicity and have direct effects on epithelial barrier integrity [17923086, 21078084].

**Immune response in AD** is facilitated essentially by immune cells that detect and process pathogens. Langerhans, IDEC, dendritic cells and macrophages, as part of the innate immune system, take up antigens penetrating through the skin barrier and stimulate Th cell differentiation [15131579, 22548207]. IL-33, produced by keratinocytes, is shown to promote proliferation, activation, and recruitment of Th2 cells, activate mast cells, ILC2 and basophils [22277940]. Th2 cells produce proinflammatory cytokines such as IL-4 and IL-13 - to reduce filaggrin in keratinocytes and disturb barrier function, IL-31 - to act on sensory nerve endings and stimulate scratching, and IL-5 - to activate eosinophils [11754819, 24373353, 14600427]. IL-4 also increases B cell IgE production [7918008, 17267416, 22430776]. IL-13, IL-4 and IL-5 are produced not only by Th2 cells, but also by ILC2 and mast cells [12145428, 7918008, 32179159, 16045735, 23603794]. Appearance of Th1, Th22 and Th17 subsets shows a transition from the acute to the chronic stage with expression of IFNG in patients with chronic lesions [28266782]. Their cytokine production (IL-17, IL-22, IL-26) also contributes to the pathogenesis of AD [27588419, 19439349, 31465744].

**In UC, immune response** maintenance is the result of impairments of tight junctions in the mucosal layer and increased uptake of luminal antigens. Activated dendritic and other antigen-presenting cells initiate the differentiation of naive T cells. Th2 produces proinflammatory cytokines IL-5, IL-4, and IL-13 [16083712, 30383225, 8757634]. IL-13, produced by NKT cells, also takes part in epithelial injury [15146247]. Th cells also differentiate into Th1, which expresses IFNG, TNF and IL-21 [20677335, 30946450]. Th17 and Treg differentiation is regulated by IL-18 [31348891] and IL-2 [30944470], which suppress inflammation in UC. Th9 cells, which are induced by IL-36 derived from the epithelium, express IL-9 and decrease cellular proliferation and weakens the intestinal barrier function [24908389]. Innate lymphoid cells ILC3s contribute to chronic intestinal inflammation and show increased cytokine production of IL-17 and IL-22 as a result of IL-23-driven inflammatory response [32332067]. Activated neutrophils form neutrophil extracellular traps [30715224]. Increased IgG level expressed by B cells is reported in UC patients [27914657].

**Pruritus in AD** is one of the main symptoms in patients with AD [30471983]. Epidermal keratinocytes can be activated by scratching, infection or irritants in response to tissue damage because of pruritus [32738956]. Various pruritogens induce itching and scratching: IL-4, IL-31, IL-13, TSLP [28890086, 24094650, 24373353, 34391816] as well as histamine produced by mast cells [35113811]. Itch is mediated by pruriceptive sensory nerve endings, which transmit the pruritic signal to the brain and result in pain [28890086]. Scratching leads to further tissue damage and activation of keratinocytes, making an inflammatory loop in AD patients [30446754].

**Chronic pain in UC** is a common symptom [22298998]. Colonic sensory neurons are sensitised by IL-13 derived from Th2 and NKT cells [36749569]. One of key mediators of neuronal excitability in UC is TNF [20427396]. Its production is elevated in UC in various immune cells and TNF receptors are expressed by neurons innervating the colon. Also serotonin signalling via HTR3 in mucosal nociceptors contribute to the visceral hypersensitivity in UC [22330338].

### Supplementary Material S4

#### Shared mechanisms in the pathway layer in key cell types of AD and UC

Below we highlight significant similarities that stand out in the AD and UC submaps. These are examples of mechanisms relevant for both AD and UC pathologies that are shown in the pathway layer.

Keratinocytes and colonocytes are the epithelial barrier cells most affected by these pathologies. They are influenced by various interleukins such as IL-13, IL-17A, IL-22, INFG through NFkB, JAK-STAT, MEK-ERK signalling pathways. Keratinocyte and colonocyte respond to the same cytokines IL-13, IL-17A, IL-22, IFNG both in AD and in UC.

Proinflammatory IL-13 is derived from Th2, ILCs, mast cells and NKT cells, acts on keratinocyte through mTOR-AKT signalling, and downregulates FLG [32922169]. Together with IL-4, it also downregulates proteins such as Loricrin, Involucrin, FLG2 that are involved in formation and maintenance of epidermis [18166499,23403047]. Similarly, in UC, IL-13 downregulates tight junction proteins Claudin8, Occludin [30619339], tricellulin [28612843] that are critical for epithelial barrier formation, and upregulates Claudin2 [16083712], which is a pore-forming protein, responsible for increased permeability for small cations. It acts through JAK-STAT pathway, as well as through IL13RA2 receptor and AP-1 activation. This way, in both pathologies IL-13 is responsible for a loss of the epithelial barrier integrity.

S100 protein activity is increased in epithelial cells in IL-17A downstream pathways through MAPKs, NF-kB signalling in AD with S100A8 and S100A9 overexpression [25308296], which decreases skin barrier function. Together with FGF2, IL-17A upregulates S100A8 through MEK-ERK in UC to contribute to mucosal barrier damage [26320657].

IL-22 signaling pathway via JAK-STAT3 contributes to epithelial barrier dysfunction by upregulating S100A8 and S100A9, and downregulating FLG expression in AD [21564072], but has a protective role in UC by upregulating tight junction protein Occludin and downregulating TNF [37440121].

In response to IFNG derived by Th1, both epithelial cells keratinocyte and colonocyte show IL-8 upregulation through JAK-STAT to attract eosinophils and monocytes in AD [32002586], and neutrophils in UC [34552702]. IL-8 is upregulated also by IL-13/IL-4 and IL-26 signalling pathways in AD keratinocyte [26147950, 31465744], and by IL-36A/IL-36G and IL-17A in UC colonocyte [26752465, 17277779].

NLRP3 inflammasome is upregulated in epithelial cells in AD and UC with increased IL-18 and IL1B production as a result [20303296, 36387340, 34133800]. Stimuli of its activation are different for AD and UC, IFNG participates in NLRP3 activation in keratinocyte [24894535], but SLC6A14 and IL-6 are involved in this activation in colonocyte [38314135, 36844140].

IL-33 acts on Th2 cells through IRAK-TRAF6-MAPKs in AD [22277940, 31455506] and in UC [25112700, 20385815] and results in influencing epithelial barrier disruption both in AD [31509236] and in UC [16083712], as well as epithelial cell differentiation [35216228, 16083712], eosinophil activation [11575456, 17620072] and nerve ending sensitisation [34391816, 36749569].

IL-4 signaling pathway in Th2 cells stimulates increased inflammatory response in both diseases. This mechanism takes place through JAK STAT6 [23303670], GATA3 [20030749] in UC and in AD [37316763, 38029848].

Th1 cells are also critical for both pathologies. IL-18, released from keratinocytes [12697661] and inflammatory dendritic epidermal cells [15131579], mediates Th1 response to enhance IFNG production through IRAK and NFkB in AD [26690141,

16723395]. Similarly, in UC, IL-18 derived from colonocytes [25736457] or dendritic cells [12920831], mediates inflammatory response acting on Th1 with increased IFNG production [12010883]. IFNG itself increases IFNG Th1 expression acting in an autocrine loop through JAK-STAT1-TBX21 as in AD [16120092] and in UC [29773813, 34751838] to maintain immune response.

These are examples of some significant mechanisms relevant for both AD and UC pathologies that are shown in the maps. A complete list of mechanisms shown in submaps for various cell types can be found in the AD and UC disease maps.

### Supplementary Material S5

Supplementary Table S5 "Complete list of overlapping groups, their content and direct links to them in corresponding disease maps" is available at https://disease-maps.io/downloads/s5.xlsx

### Supplementary Material S6

Supplementary Table S6 "Visualisation of the overlapping pathology mechanisms of UC and AD with specific connections accessible using the URLs provided" is available at https://disease-maps.io/downloads/s6.xlsx
